# K_ATP_ Channel Prodrugs Reduce Inflammatory and Neuropathic Hypersensitivity, Morphine Induced Hypersensitivity, and Precipitated Withdrawal in Mice

**DOI:** 10.1101/2022.11.10.515984

**Authors:** Alexis Doucette, Kayla Johnson, Shelby Hulke, Sunna Mujteba, Elena Miller, Peter I. Dosa, Amanda H. Klein

## Abstract

Previous studies show ATP-sensitive potassium (K_ATP_) channel openers can reduce hypersensitivity associated with chronic pain models in rodents, and reduce morphine tolerance. Many agonists of K_ATP_ channels are not soluble in physiologically relevant vehicles, requiring adaptation for clinical use. This study compared the antinociception activity of novel K_ATP_ channel targeting prodrugs, CKLP1, CKLP2, and CF3-CKLP. These prodrugs are activated by endogenous alkaline phosphatase enzymes present in the peripheral and central nervous systems. Analgesic capabilities of intrathecally injected prodrugs were tested in rodent models of spinal nerve ligation (SNL) and Complete Freund’s Adjuvant (CFA) as models for neuropathic and inflammatory pain, respectively. CKLP1 and CKLP2 significantly increased mechanical paw withdrawal thresholds 1-2 hours after intrathecal administration in the SNL model, but all three prodrugs were able to attenuate hypersensitivity up to 7 days after CFA treatment. The reduction of opioid tolerance and opioid-induced hypersensitivity in mice treated chronically with morphine was significantly reduced in CKLP1 and CKLP2 treated animals. Prodrug cleavage was confirmed in mouse spinal cords using liquid chromatography. These studies may aid in the further development of K_ATP_ channel prodrugs for use in treatments of chronic pain, opioid tolerance, and withdrawal.

## Introduction

Opioids are commonly used for the treatment of chronic pain, but long-term opioid use can lead to tolerance and eventually substance dependence. While opioids are incredibly effective therapeutics for chronic pain, it is necessary to find new therapeutics to reduce the need for opioid analgesics. Under naïve conditions, opioid medications dampen neuron excitability and neurotransmitter release, decreasing function of the nervous system (Reshef et al., 1998).ATP-sensitive potassium channels (K_ATP_ channels) are a downstream target of mu opioid receptors. Activation of K_ATP_ channels results in the hyperpolarization of neurons, preventing the propagation of action potentials and neurotransmitter release. Agonists of K_ATP_ channels are effective at reducing behavioral outputs of hypersensitivity in rodent chronic pain models (Afify et al., 2013; Niu et al., 2011). The K_ATP_ channel agonist cromakalim reduces hypersensitivity in mice when delivered intrathecally or intracerebrovascularly (Asano et al., 2000; Nakao et al., 1996). In addition, K_ATP_ channel agonists reduce opioid induced hyperalgesia and opioid tolerance in a mouse model of chronic morphine exposure (Fisher et al., 2019).

Cromakalim is not ideal for use as a therapeutic directly delivered into the nervous system or circulation due to low aqueous solubility. Formulation of prodrugs with the addition of a phosphate ester or a phosphate group with oxymethyl linker (oxymethylphosphate, OMP) significantly increases the solubility of several compounds (Rautio et al., 2008), including K_ATP_ channel agonists. Phosphate ester prodrugs are activated through dephosphorylation by the endogenously expressed enzyme alkaline phosphatase (Wiemer & Wiemer, 2015) which cleaves off the phosphate group, activating the prodrug to its parent structure. Prodrugs of cromakalim were previously designed for treatment of glaucoma by reducing intraocular pressure (Chowdhury et al., 2017). In this study we evaluate three phosphate compounds, two of which(CKLP1 and CKLP2) are prodrugs of cromakalim and one of which (CF3-CKLP1) is a prodrug of CF3-cromakalim, a more lipophilic analog of cromakalim (Roy Chowdhury et al., 2015). The high aqueous solubility of these compounds allows for easier drug delivery through intravenous, intrathecal, or subcutaneous routes. Given cromakalim’s efficacy in treating pain conditions, it was natural to question whether CKLP1, CF3-CKLP1 and/or CKLP2 share analgesic properties with the parent compound.

In this study, cromakalim prodrugs were tested for efficacy in reducing mechanical hypersensitivity in rodent models of neuropathic and inflammatory pain. The cromakalim prodrugs were also tested in mouse models of morphine tolerance and morphine precipitated withdrawal. The results showed CKLP2 and CKLP1 produced antinociception lasting 1-2 hours in a SNL model of neuropathic pain in male and female mice. Additionally, all three cromakalim prodrugs attenuated mechanical hypersensitivity in CFA-treated mice. Finally, CKLP1 and CKLP2 reduced morphine induced hypersensitivity, while CKLP2 reduced jumping and rearing behaviors after naloxone treatment. These findings support the further development of K_ATP_ channel targeting prodrugs as potential analgesic agents for treating different types of pain and reducing opioid withdrawal symptoms.

## Methods

### Animals

All experimental procedures involving animals were approved and performed in accordance with the University of Minnesota Institutional Animal Care and Use Committee guidelines. Adult male and female C57Bl6 mice were obtained from Charles River (Raleigh, NC) at five to six weeks old weighing 18.1-24.2g. Mice were acclimated to the facility on a 12-12 hour light/ dark cycle, and individual testing apparatuses prior to behavioral testing. Animals were randomly assigned to treatment groups with each group containing 5 males and 5 females. Experimenters were blinded to the drugs delivered in all behavioral experiments.

### K_ATP_ Channel Prodrug Delivery

CKLP1, CKLP2, and CF3-CKLP1 were synthesized as described (Chowdhury et al., 2016) (Figure 1). Intrathecal injections into L5 intrathecal space were performed via Hamilton syringe, PE-10 tubing and 30G dental needle set up, and delivered intrathecally as 30µg or 60µg doses in 10uL normal saline (Fairbanks, 2003).

**Figure 1.**
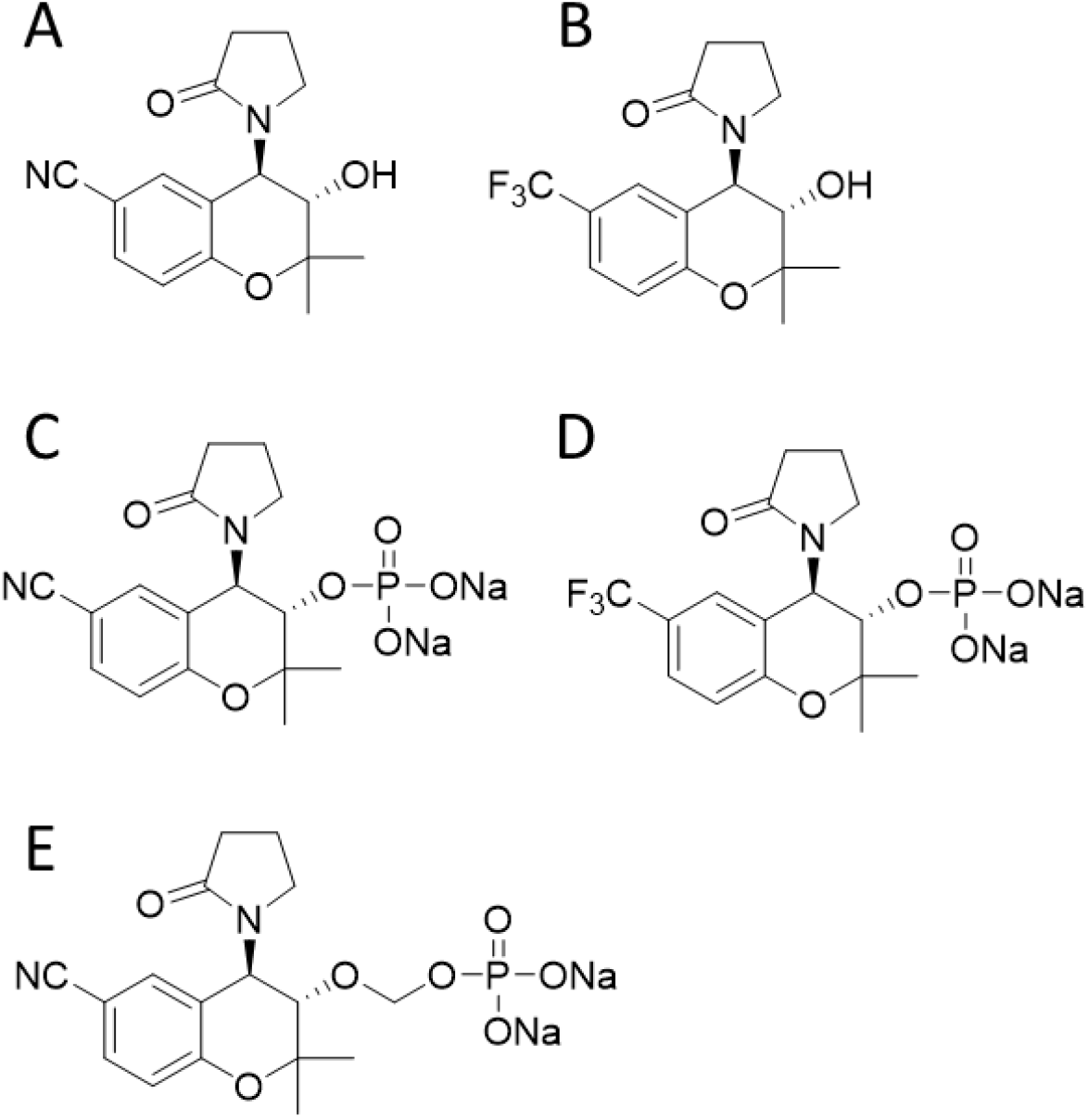
Structures of cromakalim and cromakalim prodrugs. (A) Cromakalim, (B) CF3-Cromakalim, (C) CKLP1, (D) CF3-CKLP1, and (E) CKLP2

### Mechanical Paw Withdrawal Thresholds

Animals were placed in testing apparatus for acclimation on two separate occasions lasting 45-60 minutes. The testing apparatus consisted of individual acrylic chambers on a mesh floor to allow access to paws for testing. Mechanical paw withdrawal thresholds were measured using electronic von Frey testing equipment (Electric von Frey Anesthesiometer, 2390, Almemo ® 2450, IITC Life Science, Woodland Hills, CA). Hind paw plantar surfaces were gently pressed with the probe until a nocifensive response was observed. Baseline measurements were collected five times per hind paw and averaged. Single measurements were used for post-drug SNL time-course experiments. An average of 5 measurements per paw were used for mechanical testing in the post-drug CFA model, and 3 measurements per paw were used under opioid tolerance tests.

### Spinal Nerve Ligation

A spinal nerve ligation model of neuropathic pain was used to determine the effective time-course of cromakalim prodrugs as analgesics (Rigaud et al., 2008). Mice were anesthetized with isoflurane in oxygen, a section of the L4 spinal nerve was removed from the animal (Rigaud et al., 2008; Ye et al., 2015), and the surgical site was closed with absorbable sutures. After healing for 14 days, mechanical paw withdrawal thresholds were collected on the ipsilateral and contralateral hind paws 30 min, 1 hour, 1.5 hours, 2 hours, 4 hours, 6 hours, and 8 hours post intrathecal injection of K_ATP_ channel prodrugs.

### Complete Freund’s Adjuvant

To test the efficacy of cromakalim prodrugs to reduce hypersensitivity in a model of inflammatory pain, 5 male and 5 female mice received a unilateral intraplantar injection of CFA (10uL, F5881, Sigma Chemical, St. Louis, MO). Mechanical paw withdrawal thresholds were collected prior to CFA injection, and one-hour post intrathecal injection CKLP1, CF3-CKLP1, CKLP2, and saline (30ug and 60ug in 10uL). Further mechanical paw withdrawal threshold measurements were taken on days 2, 3 and 7 post CFA administration.

### Morphine Tolerance

Morphine tolerance was established in naïve mice via twice daily injections for 5 days (∼0800hrs and 1700hrs, 15mg/kg. s.c., 100µl, Sigma Chemical, St. Louis, MO) (Liang et al., 2011). Animals received intrathecal injection of prodrug (30 µg or 60uµg in 10ul saline) or saline control 30 min prior to receiving subcutaneous morphine. Mechanical paw withdrawal testing was performed before administration of morphine and 30 minutes after morphine.

### Naloxone Precipitated Withdrawal

On day 6 after morphine tolerance was completed, mice were given final subcutaneous dose of morphine (15mg/kg in 100uL saline) and were placed in 10cm by 15cm Plexiglas chambers.

Animal behavior was video recorded for 15 minutes to determine baseline animal behavior, 30 minutes after morphine administration. Two hours post-morphine, intrathecal injection of CKLP1, CKLP2, CF3-CKLP1, or saline control (60µg in 10µL, i.t.) was administered.

Withdrawal was precipitated with intraperitoneal injection of naloxone hydrochloride (1mg/kg in 100uL saline). Immediately after naloxone injection, animal behavior was video recorded for 15 minutes. The number of jumps and bouts of rearing behaviors were counted in one-minute intervals by two independent scorers (El-kadi & Sharif, 1994).

### Tissue Collection and Tissue Sample Preparation

Animals used in CFA and morphine tolerance and withdrawal experiments were euthanized 24 hours after final injection of prodrug. Spinal cords were harvested from animals and immediately frozen at -20°C and stored until used for conversion analysis and prepared as described previously (Nakhi et al., 2021). Animal tissue was transferred to a clean microcentrifuge tube prechilled with 200ul of mobile phase starting solution (2.5mM ammonium formate in 5% acetonitrile pH 2.15). Tissue was homogenized on ice using 1.5 mL micro-tube sample pestles until no tissue remnants were visible. The pestle was rinsed with an additional 100ul of mobile phase mixture. Homogenate was vortexed and placed in prechilled (4°C) centrifuge for 25 minutes at 16,100rcf. Supernatant was transferred to a clean tube. 120ul of homogenate was mixed with 5ul of flavopiridol internal standard and placed into amber autosampler vial with 200ul insert.

### Liquid Chromatography

A Thermo Scientific Ultimate 3000 UHPLC+ focused pump, autosampler and column compartment were used with Chromeleon 7.2 software to collect liquid chromatography data. A Waters XBridge BEH C18 130Å, 3.5um, 3mm x 50mm column was maintained at 30°C. Biological samples were stored at 4°C when not injecting to suppress additional enzymatic activity prior to *ex vivo* analysis. Injection volumes of 2ul were used. Mobile phase A consisted of 2.5mM ammonium formate in water at a pH of 2.15 and mobile phase B was 2.5mM ammonium formate in acetonitrile. The column was preequilibrated from 10/90 Mobile A/B to 95/5 Mobile A/B over 2 minutes, then held at 95/5 Mobile A/B for one minute prior to injection. 95/5 Mobile A/B was held for 3 minutes after injection. The mobile phases were then ramped to 80% Mobile B over 10 minutes for elution of prodrug and cromakalim and held at 20/80 mobile A/B for 5 minutes. The method used a constant flow rate of 0.8mL/min while monitoring absorbance of 252nm for prodrugs, cromakalim, flavopiridol internal standard and other products. Five-point standard curves of CKLP1, CKLP2 and cromakalim (1.563uM to 100uM) were made in extraction buffer for quantification.

### Data Analysis

Animal data was collected and analyzed by blinded personnel. ANOVA (one-way and two-way repeated measures) with Dunnett’s *post hoc* analysis were used to determine significance for mechanical thresholds and withdrawal behaviors. GraphPad Prism version 9 (GraphPad Software, San Diego, CA) was used for statistical analysis. HPLC chromatograms were automatically integrated with Chromeleon 7.2, and manually checked for consistency.

Data presented as mean ± S.D.

## Results

### K_ATP_ channel targeting prodrugs increase mechanical withdrawal thresholds in a mouse model of neuropathic pain

To determine the efficacy of cromakalim prodrugs against neuropathic pain, mechanical paw withdrawal thresholds were tested 14 days after SNL. A significant increase in mechanical withdrawal thresholds on the ipsilateral hind paw were found with the 60ug dose of CKLP1 at 30 minutes, and the 30ug dose had a significant increase at 480 minutes after intrathecal administration when compared to saline treated animals (Figure 2a). CKLP2 increased the ipsilateral hind paw thresholds with the 60µg dose at 60 minutes and 90 minutes post intrathecal injection compared to saline (Figure 2b). CF3-CKLP1 significantly increased the mechanical paw withdrawal thresholds at the 30µg dose at 360 minutes and 480 minutes compared to saline (Figure 2c). After normalizing to baseline for each individual treatment group, the area under the curve for CKLP1 and CKLP2 were significantly increased over saline treated animals 2 hours after injection, as was the 30µg dose of CF3-CKLP1 (Figure 2d). The 60-minute post injection time point was chosen for further experiments due to the increase in mechanical paw withdrawal thresholds for multiple prodrug treatment groups and dosages.

**Figure 2.**
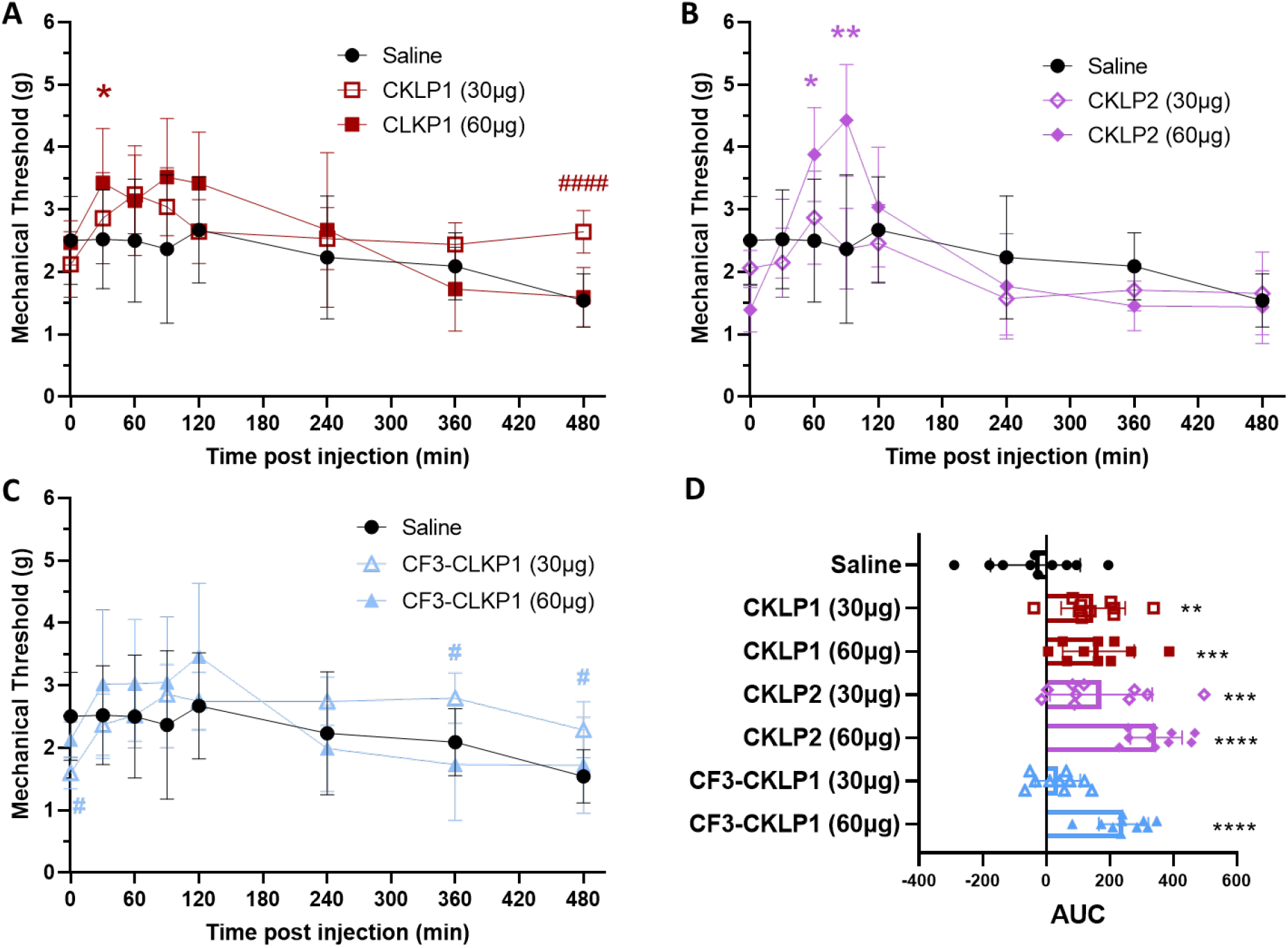
Intrathecal administration of CKLP1, CKLP2, and CF3-CKLP1 increased mechanical paw withdrawal thresholds in spinal nerve ligated mice. (A) CKLP1 significantly increased paw withdrawal thresholds compared to saline treated animals 30 minutes post injection at the 60ug dose on the ipsilateral hind paw (Repeated measures ANOVA, Time x Drug effect F(14, 189) = 2.867, *P* = 0.0006). The 30ug dose of CKLP1 was significantly higher than saline at 480 minutes in the ipsilateral hind paw. (B) CKLP2 at a 60ug dose was significantly increased compared to saline at 60 and 90 minutes (repeated measures ANOVA, Time x Drug effect F(14, 189) = 6.366, *P* < 0.0001). (C) CF3-CKLP1 at a 30ug dose was significantly higher than saline at 360 minutes and at 480 minutes (repeated measures ANOVA, Time x Drug effect F(14, 189) = 2.995, *P* =0.0004). Data is plotted as the treatment group average with SD, #, #### indicates significance of 30ug dose with P <0.05, and <0.001, respectively, *, ** indicates significance of 60ug dose at P<0.05 and 0.01, respectively. (D) Area under the curve for data points within the first 2 hours of drug administration. **, ***, **** indicate significnce at P< 0.01, 0.001, and 0.0001 compared to saline injection, respectively. n=10 with 5 male and 5 female mice per group.

### K_ATP_ channel targeting prodrugs reduce mechanical hypersensitivity in a mouse model of inflammatory pain

Mice were injected with CFA in one hind paw and mechanical thresholds were measured at 70 minutes, as well as 24, 36, and 168 hours after CFA injection. The 30µg and 60µg doses of CKLP1 significantly increased mechanical paw withdrawal thresholds at all time points tested post-CFA injection compared to saline (Figure 3a). The 30µg dose of CKLP2 significantly increased mechanical paw withdrawal thresholds at 24 hours and 168 hours post CFA injection, and the 60µg dose of CKLP2 significantly increased thresholds at one hour, 24, 72 and 168 hours compared to saline (Figure 3b). The 30µg dose of CF3-CKLP1 significantly increased mechanical paw withdrawal thresholds at all time points tested post CFA injection compared to saline (Figure 3c). The 60µg dose of CF3-CKLP1 significantly increased thresholds in the ipsilateral hind paw at 24, 48 and 168 hours post CFA injection when compared to saline (Figure 3c).

**Figure 3.**
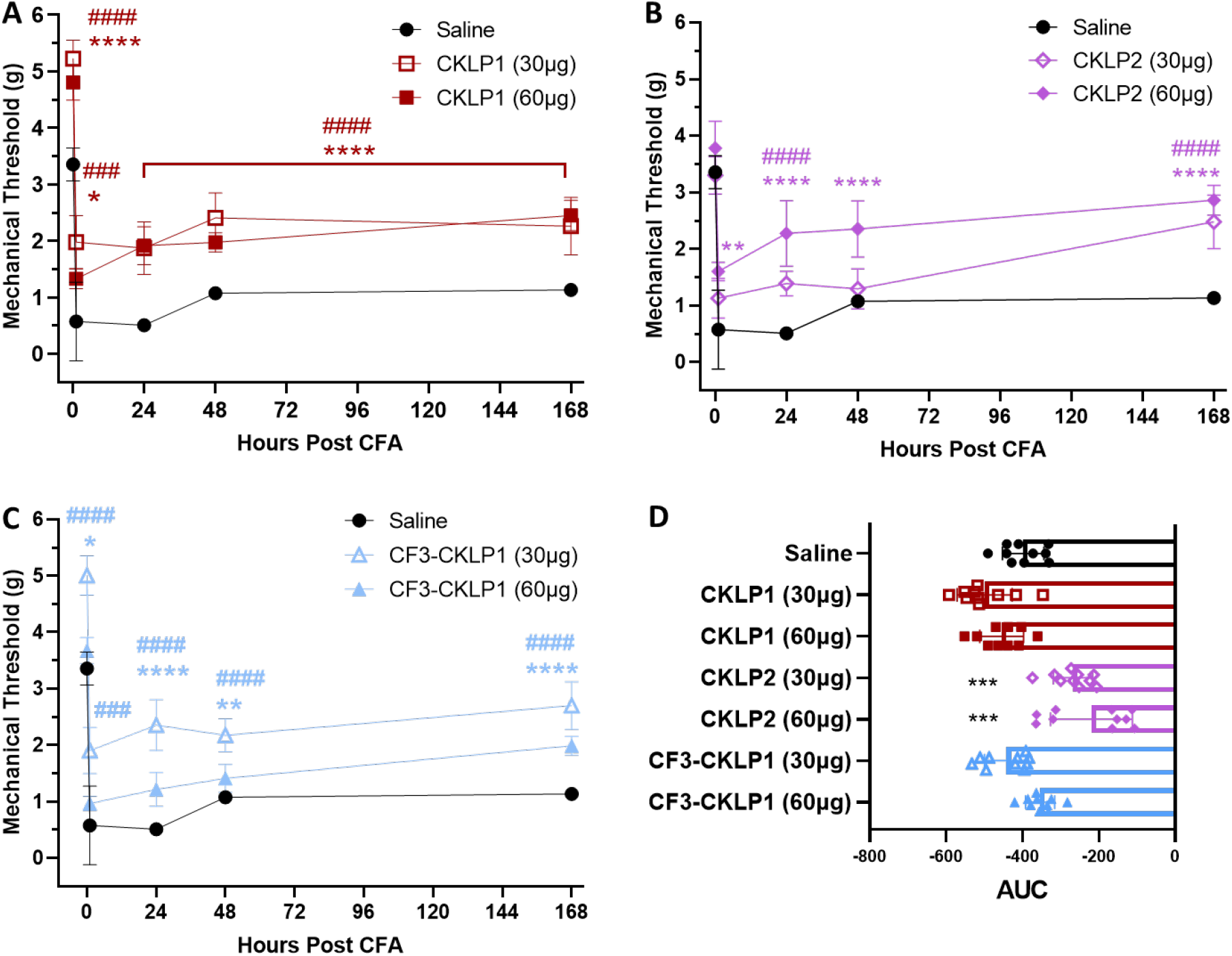
Intrathecal administration of cromakalim prodrugs increased mechanical paw withdrawal thresholds in mice treated with Complete Freund’s Adjuvant. (A) In the ipsilateral hind paw, the 60ug dose of CKLP1 increased mechanical paw withdrawal thresholds on days 1, 2, 3 and 7 after CFA injection (Repeated measures ANOVA, Time x Drug effect F(8, 108) = 4.749, *P* < 0.0001). The 30ug dose of CKLP1 increased mechanical paw withdrawal thresholds on days 1, 2, 3 and 8 post CFA in the ipsilateral hind paw. (B) The 30ug dose of CKLP2 increased mechanical paw withdrawal thresholds on day 2 and day 8 (Repeated measures ANOVA, Time x Drug effect F(8, 108) = 9.555, *P* < 0.0001). The 60ug dose of CKLP2 increased mechanical paw withdrawal thresholds on days 1, 2, 3, and 8. (C) The 30ug dose of CF3-CKLP1 increased mechanical paw withdrawal thresholds on all days (Repeated measures ANOVA, Time x Drug effect F(8, 108)=3.684, *P* =0.0008). The 60ug dose of CF3-CKLP1 increased mechanical paw withdrawal thresholds on days 2, 3 and 8 in the ipsilateral hind paw. Data is plotted as the treatment group average with SD, #, ###, #### indicates significance of 30ug dose with P <0.05, <0.001, and <0.0001 respectively. *, **, **** indicates significance of 60ug dose at P<0.05, 0.01, and <0.0001 respectively compared to saline treatment group. n=10 with 5 male and 5 female mice per group.

### Opioid induced hypersensitivity is mitigated by CLKP1, CKLP2, and CF3-CKLP1

Morphine tolerance was established with twice daily morphine injections (15mg/kg) for five days with prodrug injections (60µg in 10ul saline) occurring one-hour prior to morphine injection. In order to measure the effects of K_ATP_ channel prodrugs on opioid induced hyperalgesia (OIH) and morphine tolerance, mechanical paw withdrawal thresholds were measured pre-and-post morphine, respectively. CKLP1 and CKLP2 increased mechanical paw withdrawal thresholds pre-morphine and post-morphine when compared to saline starting on Day 2 and Day 4, respectively (Figure 4).

**Figure 4.**
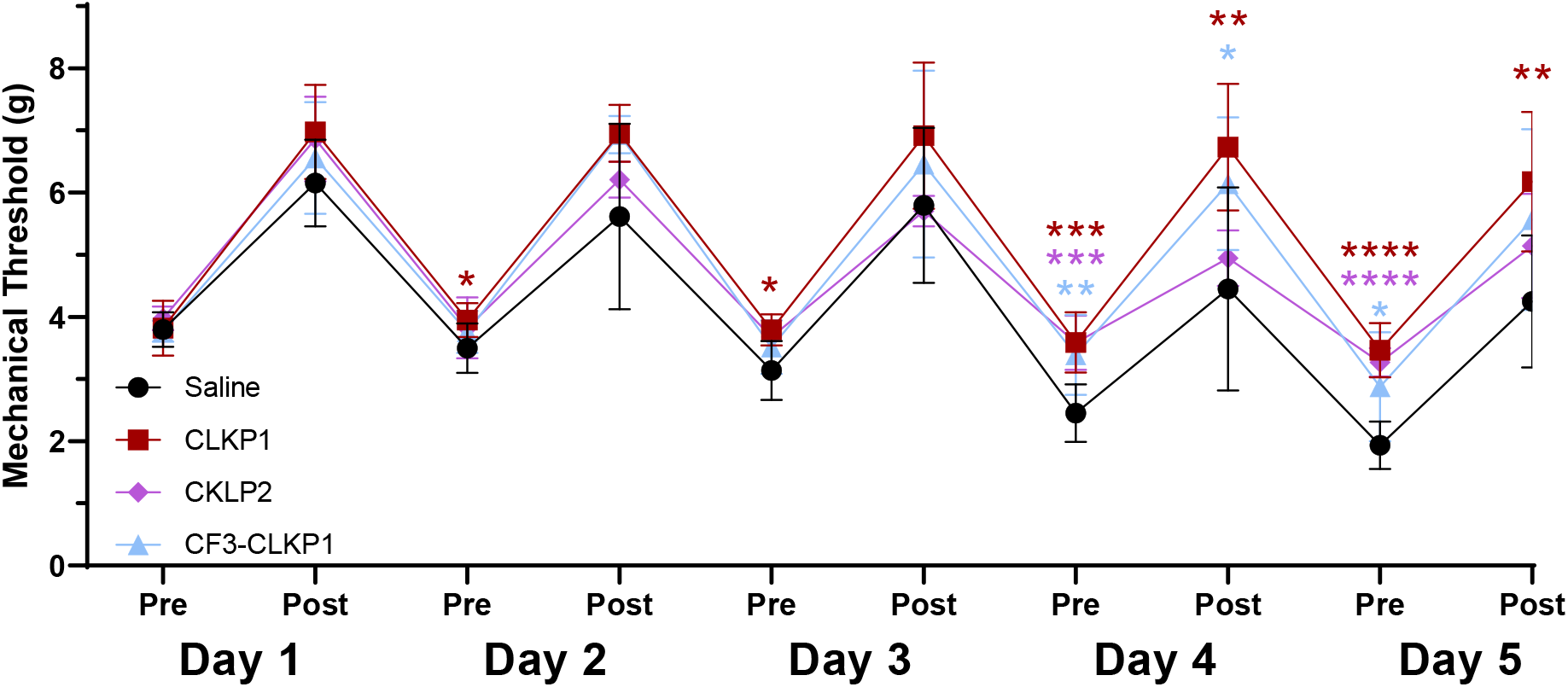
Cromakalim prodrugs attenuate opioid induced hyperalgesia and opioid tolerance. Morphine was delivered twice daily for five days (15 mg/kg, sc) and CKLP1, CKLP2, CF3-CKLP1 (60μg, it) or vehicle was administered 30 minutes before morphine. A significant increase in paw withdrawal thresholds in CKLP1 and CKLP2 treated mice were seen pre-and post-morphine administration, largely on days 3-5 (Repeated measures ANOVA, Drug effect: F (3, 35) = 7.423). Mechanical thresholds post-morphine were collected 30 minutes after injection. Data plotted as the treatment group average with SD, \**P* <0.05, \*\**P* <0.01, \*\**\*P* <0.001, \*\*\*\**P* <0.0001. n = 5 male and 5 female mice per group.

### Precipitated withdrawal behaviors are attenuated by CLKP2

Withdrawal in mice was induced with naloxone following five days of twice daily morphine and prodrug injection (see above). The total number of jumps and rearing bouts in the first five minutes of withdrawal were scored for each animal and averaged within the treatment group. No significant changes in the number of jumps or rearing were observed in animals treated with CKLP1 or CF3-CKLP1 compared to the saline treated animals (Figure 5a and 5b). A significantly lower number of jumps and rearing bouts were observed in mice treated with CKLP2 during the first five minutes after naloxone administration (Figure 5a and 5b).

**Figure 5.**
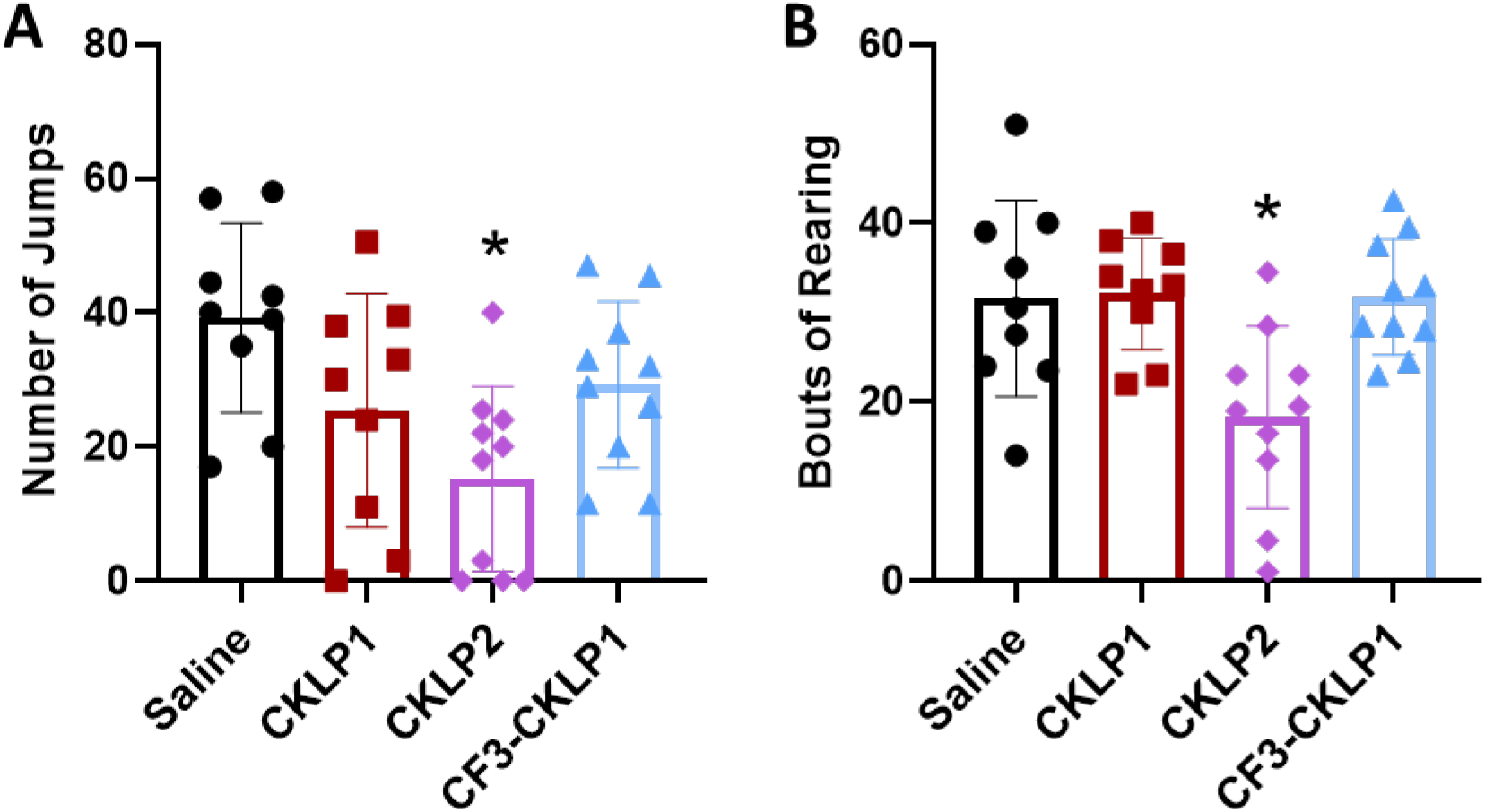
CKLP2 decreases the number of jumps and rearing bouts in mice after precipitated morphine withdrawal. Two hours post-morphine (15 mg/kg, sc) and one hour post intrathecal injection of CKLP1, CKLP2, CF3-CKLP1, or saline control (60µg, i.t.) withdrawal was precipitated with intraperitoneal injection of naloxone hydrochloride (1mg/kg in 100uL saline). (A) CKLP2 decreased withdrawal jumps compared to saline (One-way ANOVA, F(3, 34) = 4.455). (B) CKLP2 also decreased withdrawal rearing compared to saline (One-Way ANOVA, F(3, 34) = 4.455). CKLP1 and CF3-CKLP1 did not have a significant effect on jumping or rearing when compared to saline. Data is expressed as the total number of behaviors per animal in 5 minutes averaged for each treatment group ± SD. \**P* <0.05, n= 5 males and 5 females per group.

### HPLC Analysis

HPLC was used to separate cromakalim, CKLP1, CKLP2, and flavopiridol with retention times of 7.82 ±0.03 min, 6.50 ±0.04 min, 6.85 ±0.01 min, and 7.99 ±0.04 min, respectively.

Compound concentrations were calculated from standard curves plotted as peak area of interest compared to internal standard peak area versus the compound concentration. We were unable to determine detectable levels of CKLP1 or CKLP2 in spinal cord samples from animals receiving intrathecal injection of either compound. Cromakalim was detected in three of the 30 animals treated with CKLP1. The average concentration of cromakalim was 2.49± 0.45uM in animals intrathecally injected with 60ug of CKLP1. Cromakalim was detected in 11 of the 30 animals treated with CKLP2. The average concentration of cromakalim was 2.22± 0.31uM in animals injected with 60ug (n = 8) and 1.17± 0.11uM in animals given 30ug CKLP2 (n = 3). CF3-CKLP1 was not able to be quantified with reverse phase HPLC because the compound eluted in the solvent front. We were unable to establish analytical conditions for the unambiguous quantification of CF3-CKLP1 so the concentration of this compound was not determined. No compounds were detected in saline treated animals.

## Discussion

Cromakalim prodrugs effectively increased the mechanical paw withdrawal thresholds in mouse models of chronic neuropathic and inflammatory pain. The prodrugs tested also increased mechanical paw withdraw thresholds under chronic morphine pre- and post-morphine, as well as decreased opioid withdrawal behaviors. This study is novel because the analgesic efficacy, tolerance attenuation, and pharmacokinetic properties and conversion of the cromakalim prodrugs systemically *in vivo*, especially in mouse spinal cord, were relatively unknown. We conclude that future iterations of K_ATP_ channel targeting prodrugs are worthy of investigation in the future.

Cromakalim has previously been demonstrated to have analgesic capabilities, similar to other K_ATP_ channel agonists in chronic pain models in rodents (Du et al., 2011; Luu et al., 2019; Niu et al., 2011; Wu et al., 2011). The cromakalim prodrugs CKLP1, and CKLP2 were effective at increasing mechanical paw withdrawal in a sciatic nerve ligation model of chronic neuropathic pain and a CFA model of inflammatory pain in mice. The expression of alkaline phosphatase increases under nerve injury (Brichacek & Brown, 2019), chronic inflammation conditions (Park et al., 2020), and gastrointestinal inflammation (Fawley & Gourlay, 2016), which is in alignment with the idea that the cromakalim prodrugs could be effective as analgesics. The time frame from injection to the maximum paw withdrawal latency (∼60-90 min) during the SNL model is likely due to the timing of alkaline phosphatase dephosphorylation of the prodrugs, converting them to the parent compound cromakalim. Previous pharmacokinetic analysis of CKLP1 indicates that these compounds have a plasma half-life up to 5 hours (Roy Chowdhury et al., 2021), which explains why some of the compounds have continued efficacy several hours after injection.

In contrast, the cromakalim prodrug CF3-CKLP1 increased mechanical paw withdrawal thresholds only in the CFA mouse model of inflammatory pain, similar to CKLP1 and CKLP2. K_ATP_ channel agonists have been shown to have anti-inflammatory properties, possibly due to inhibiting cytokine release from microglia (Plachinta et al., 2004; Virgili et al., 2011). This could explain why all three cromakalim prodrugs would increase the mechanical paw withdrawal thresholds in the inflammatory pain model, but not in the neuropathic pain model. In addition, cromakalim is a known vasodilator, which may increase local blood flow at lumbar spinal segments after intrathecal injection, inhibiting pain in a similar mechanism to spinal cord stimulation (Wu et al., 2008). Afferent fibers containing TRPV1 are necessary for vasodilation to occur during spinal cord stimulation, which is partially mediated by extracellular signal-regulated kinase (ERK)(Wu et al., 2006). K_ATP_ channel agonists increase ERK phosphorylation in vitro, but do not affect total ERK concentrations. Therefore, it is possible that opening K_ATP_channels could have multiple mechanisms of producing analgesia during inflammation when delivered intrathecally.

Cromakalim prodrugs were able to decrease opioid induced hyperalgesia during chronic morphine exposure. The greatest effect was seen with CKLP1 and CKLP2 treatment, as mechanical paw withdrawal threshold values were increased on days four and five of chronic morphine exposure compared to vehicle treated animals. CF3-CKLP1 also decreased opioid induced hyperalgesia, but to a smaller extent. The pairing of CKLP1 with morphine also reduced opioid tolerance as seen by measurements obtained post-morphine administration compared to the saline treated animals. A small increase in mechanical thresholds was observed with CKLP2 treatment on day five of chronic morphine exposure. A large decrease in withdrawal behaviors was seen in CKPL2 treated mice, including a decreased number of jumps and rearing bouts compared to vehicle control animals. The decrease in morphine-induced hyperalgesia and withdrawal symptoms might be due to the opening of K_ATP_ channels independently of opioid signaling, decreasing overall neuron excitability (Yang et al., 2014). Systemic delivery of K_ATP_ channel openers, including cromakalim, decrease symptoms of withdrawal, while K_ATP_ channel blockers increase withdrawal symptoms in rodent models (Robles et al., 1994; Seth et al., 2010). Symptoms of withdrawal intensify under increased concentrations of ATP in the central nervous system (Burma et al., 2017) and blocking pannexin 1 channels, and subsequently ATP-release, has been shown to alleviate withdrawal symptoms in rodents(Burma et al., 2017). K_ATP_ channel agonists could counteract increased concentrations of ATP which naturally inhibit K_ATP_ channels, therefore decreasing withdrawal symptoms.

The HPLC data demonstrate the phosphate group is cleaved after intrathecal injection from the cromakalim prodrugs, converting the prodrug into the cromakalim parent compound.Cromakalim was present in animals treated with CKLP1 and CKLP2 in the spinal cord 24-hours after injection. We believe the antinociceptive responses seen in the behavioral data are attributed to the presence of cromakalim in the spinal cord of these animals. The low concentrations of cromakalim in prodrug treated animal samples could be from low extraction efficiencies, or elimination of the compounds from the spinal cord over a 24-hour period of time. Differences in alkaline phosphatase expression or activity may occur under different injury models, so it is possible that conversion of prodrug into cromakalim would occur at different rates between models. A direct comparison between, for example, inflammatory and neuropathic pain conditions and alkaline phosphatase activity in the peripheral or central nervous system has not been done to our knowledge.

Levels of alkaline phosphatase have been shown to increase in the spinal column under nerve injury state in humans (Citak et al., 2016; Vimalraj, 2020) and the peripheral nervous system in rodents(Pinner & Campbell, 1965). Due to the increased expression and/or activity of alkaline phosphatase during injury conditions, we hypothesized that conversion of prodrugs would occur under painful syndromes and predicted compounds with greater phosphate group cleaving would have higher analgesic efficacy. The overall success of CKLP2 in all experiments is likely attributed to the additional carbon and oxygen linking the phosphate group to cromakalim. This type of OMP-prodrug is known to more efficiently activated in vivo than compounds in which the phosphate group is directly attached to an alcohol (e.g. CKLP1)(Rautio et al., 2008).

Due to the overall efficacy of CKLP2 as an analgesic in neuropathic and inflammatory pain, as well as decreasing opioid tolerance and withdrawal symptoms, it would be beneficial to move forward with investigating CKLP2 or investigating similar analogues for other K_ATP_channel openers. K_ATP_ channel agonists more specific to SUR1 subtypes most highly expressed in the peripheral nervous system and spinal cord would be worth investigating (Fagerberg et al., 2014; Moreau et al., 2000). Other K_ATP_ channel openers more specific to SUR2 subtypes, such as pinacidil, show antinociception capabilities in rodent models, but would be expected to have cardiovascular side effects if introduced systemically due to activity on cardiovascular and endothelial cells (Luu et al., 2019; Moreau et al., 2000).

In conclusion, these data indicate K_ATP_ targeting prodrugs can produce antinociception and analgesia in animals. CKLP2 likely initiates this effect through rapid dephosphorylation which is sustained after several hours. In addition, CKLP1 and CKLP2 attenuated morphine induced hypersensitivity, and CKLP2 reduced mouse behaviors associated with morphine after chronic morphine administration. Further investigation into K_ATP_ channel targeting prodrugs and their antinociceptive effects and exploration of their efficacy in different pain conditions, with or without opioid treatment, may aid toward developing novel therapeutics for treating pain conditions and reducing overall consumption of opioids as analgesics for chronic pain.

## Acknowledgments

We thank Dr. Melissa Maurer-Jones of the UMD Department of Biochemistry and Chemistry for help with liquid chromatography.

## Authorship Contributions

*Participated in research design:* Klein and Dosa.

*Conducted experiments*: Doucette, Johnson, Hulke, Mujteba, Miller.

*Performed data analysis*: Doucette and Klein

*Wrote or contributed to the writing of the manuscript:* Doucette, Johnson, Dosa, and Klein

